# Proteolysis and neurogenesis modulated by LNR domain proteins explosion support male differentiation in the crustacean *Oithona nana*

**DOI:** 10.1101/818179

**Authors:** Kevin Sugier, Romuald Laso-Jadart, Soheib Kerbache, Jos Kafer, Majda Arif, Laurie Bertrand, Karine Labadie, Nathalie Martins, Celine Orvain, Emmanuelle Petit, Julie Poulain, Patrick Wincker, Jean-Louis Jamet, Adriana Alberti, Mohammed-Amin Madoui

## Abstract

Copepods are the most numerous animals and play an essential role in the marine trophic web and biogeochemical cycles. The genus *Oithona* is described as having the highest numerical density, as the most cosmopolite copepod and iteroparous. The *Oithona* male paradox obliges it to alternate feeding (immobile) and mating (mobile) phases. As the molecular basis of this trade-off is unknown, we investigated this sexual dimorphism at the molecular level by integrating genomic, transcriptomic and protein-protein interaction analyses.

While a ZW sex-determination system was predicted in *O. nana*, a fifteen-year time-series in the Toulon Little Bay showed a biased sex ratio toward females (male / female ratio < 0.15±0.11) highlighting a higher mortality in male. Here, the transcriptomic analysis of the five different developmental stages showed enrichment of Lin12-Notch Repeat (LNR) domains-containing proteins coding genes (LDPGs) in male transcripts. The male also showed enrichment in transcripts involved in proteolysis, nervous system development, synapse assembly and functioning and also amino acid conversion to glutamate. Moreover, several male down-regulated genes were involved in the increase of food uptake and digestion. The formation of LDP complexes was detected by yeast two-hybrid, with interactions involving proteases, extracellular matrix proteins and neurogenesis related proteins.

Together, these results suggest that the *O. nana* male hypermotility is sustained by LDP-modulated proteolysis allowing the releases and conversions of amino acid into the excitatory neurotransmitter glutamate. This process could permit new axons and dendrites formation suggesting a sexual nervous system dimorphism. This could support the hypothesis of a sacrificial behaviour in males at the metabolic level.

## Introduction

Copepods are small planktonic crustacean forming the most abundant metazoan subclass on Earth and occupy all ecological aquatic niches (Huys and Boxshall 1991; Kiørboe 2011). Among them, the genus *Oithona* is described as having the highest numerical density (Gallienne and Robins 2001), cosmopolite (Nishida 1985) and playing a key role of secondary producer in the marine food web and biogeochemical cycles (Steinberg and Landry 2017). Because of its importance, *Oithona* phylogeography, ecology, behaviour, life cycle, anatomy and genomics are studied (Cornils, Wend-Heckmann, and Held 2017; Dvoretsky and Dvoretsky 2009; Kiørboe 2007; Madoui et al. 2017; Mironova and Pasternak 2017; Paffenhöfer 1993; Sugier et al. 2018; Zamora-Terol et al. 2014; Zamora Terol 2013).

*Oithona* is an ambush feeder: to feed, the individual remains static, jumps on preys that come on its range, and captures them with its buccal appendages (Kiørboe 2007). While females are mostly feeding and thus static, males actively seek females for mating. The male mating success increases by being motile and non-feeding, but its searching activity is limited by the energy resources previously stored. Theoretically, to maximise mating success, males have to alternate feeding and female searching periods which constitutes a paradox in the *Oithona* male behaviour (Kiørboe 2007).

In the Little Bay of Toulon, *O. nana* is the dominant zooplankton throughout the year without significant seasonal variation suggesting a continuous reproduction (Richard and Jamet 2001) as observed in others *Oithona* populations (Temperoni et al. 2011). Under laboratory condition with two sexes incubated separately, *O. nana* males have a mean lifetime of 25 days and 42 days for females. However, the lifespan *in situ* is unknown as well as its reproduction rate. Nonetheless, in the case of female saturation, *O. davisae* males have a reproduction rate (0.9 females male^−1^ day^−1^) depending on the production of spermatophores that are transferred during a mating (Kiørboe 2007).

A biased sex-ratio toward females (male/female ratio<0.22) was observed in *O. nana* population of the Toulon Little Bay (Richard and Jamet 2001). Several causes could explain this observation. One factor evoked was the higher male exposure to predators due to its higher motility; this behaviour was assimilated to a “risky” behaviour (Hirst et al. 2010; Kiørboe 2006). However, we can propose other possibilities such as environmental sex determination (ESD) that has already been observed in other copepods (Voordouw and Anholt 2002) but also energy resource depletion as a consequence of male energy consumption during mate search. A risky behaviour implies environmental factors decreasing fitness, like encountering a predator. In contrast, a sacrificial behaviour is deterministic and thus only driven by genetic factors, for example, forcing the consumption of all energy resources to find a partner or die while trying. The sacrifice of males is observed in semelparous animals (single reproductive cycle in a lifetime) like insects or arachnids and is associated to female fitness increase. This behaviour has never been described in iteroparous animals like copepods (multiple reproductive cycles in a lifetime) (Hairston and Bohonak 1998).

Recently, the *O. nana* genome was sequenced, and its comparison to other genomes showed an explosion of Lin-12 Notch Repeat (LNR) domains-containing proteins coding genes (LDPGs) (Madoui et al. 2017). Among the 75 LDPGs present in the genome, five were found under natural selection in Mediterranean Sea populations, including notably one-point mutation generating an amino-acid change within the LNR domain of a male-specific protein (Arif et al. 2019; Madoui et al. 2017). This provided a first evidence of *O. nana* molecular differences between sexes at the transcriptional level and a potential gene repertoire of interest.

To further investigate the molecular basis of *O. nana* sexual differentiation, we propose in this study a multi-approach analysis including, (i) *in situ* sex ratio determination in time series (ii) sexual system determination by sex-specific polymorphism analysis; (iii) *in silico* analysis of the structure and evolution of the LDPGs, (iv) sex-specific gene expression through RNA-seq analysis and (v) LDPs interaction protein network by yeast two-hybrid.

## Material and methods

### Sex-ratio report in the Toulon Little Bay

*Oithona nana* specimens were sampled at the East of the Toulon Little Bay, France (Lat 43° 06’ 52.1” N and Long 05° 55’ 42.7” E). The samples were collected from the upper water layer (0-10m) using zooplankton nets with a mesh of 90µm and 200µm. Samples were preserved in 5% formaldehyde. The monitoring of *O. nana* in Toulon Little Bay was performed from 2002 to 2017. Individuals of both sexes were identified and counted under the stereomicroscope.

### Biological materials and RNA-seq experiments

For the RNA-seq experiment, the plankton sampling took place in November 2015 and November 2016 using the same collecting method than previously described. The samples were preserved in 70% ethanol and stored at −20°C. The *Oithona nana* were isolated under the stereomicroscope. We selected individuals from five different development stages: five pairs of egg-sac, four nauplii (larvae), four copepodites (juveniles), four female adults and four male adults. All individuals were isolated from the November 2015 sample, except for eggs. Each individual was transferred alone and crushed with a tissue grinder (Axygen) into a 1.5 mL tube (Eppendorf). Total mRNAs were extracted with the NucleoSpin RNA XS kit (Macherey-Nagel) following the manufacturer instructions, then quantified on Qubit 2.0 with the RNA HS Assay kit (ThermoFisher Scientific) and quality assessed on Bioanalyzer 2100 with the RNA 6000 Pico Assay kit (Agilent). cDNAs were constructed using the SMARTer v4 Ultra low Input RNA kit (Takara). After cDNA shearing by Covaris E210 instrument, Illumina libraries were constructed using the NEBNext Ultra II kit (New England Biolabs) and sequenced on Illumina HiSeq2500. A minimum of 9.7e^6^ reads pairs was produced from each individual (Supplementary Notes S1).

### Sex-determination system identification by RNA-seq

RNA-seq reads of both sexes (four females and four males) were alignment against predicted cDNA. Reads having an alignment length ≤ 80% and identity cut-off ≤ 97% were removed. The variant calling step was performed with the ‘*samtools mpileup*’ and ‘*bcftools call*’ commands, with default parameters (Li et al. 2009) and only bi-allelic sites were kept.

To identify the most likely sexual system in *O. nana*, we used *SD-pop* (Käfer, Lartillot, Picard & Marais, *in prep*). Just like its predecessor *SEX-DETector* (Muyle et al. 2016), *SD-pop* calculates the likelihood of three sexual models (absence of sex chromosomes, XY system or ZW system) which can be compared using their BIC (Bayesian Information Criterion). The difference with *SEX-DETector* is that *SD-pop* is based on population genetics (i.e. Hardy-Weinberg equilibrium for autosomal genes, and different equilibria for sex-linked genes) instead of mendelian transmission from parents to offspring, and thus can be used without the requirement of obtaining a controlled cross.

The number of individuals used (four for each sex) is close to the lower limit for the use of *SD-pop*, where the robustness of the method is weakening. To test whether the model preferred by *SD-pop* could have been preferred purely by chance, we permuted the sex of the individuals, with the constraint of keeping four females and four males ((8!/(4!*4!)-1=69 permuted datasets). As the XY model is strictly equivalent to the ZW model with the sexes of all individuals changed, two *SD-pop* models (no sex chromosomes, ZW) were run on all possible permutations of the data, and the BIC of each model was calculated. The genes inferred as sex-linked based on their posterior probability (>0.8) were manually annotated.

### *Arthropoda* phylogenetic tree

The ribosomal 18S sequences from seven arthropods including five copepods (*O. nana, Lepeophtheirus salmonis, Tigriopus californicus, Eurytemora affinis, Calanus glacialis, Daphnia pulex and Drosophila melanogaster*) were downloaded from NCBI. The sequences were aligned with MAFFT (Katoh and Standley 2013) using default parameters. The nucleotide blocks conserved in the seven species were selected by Gblock on Seaview (Gouy, Guindon, and Gascuel 2010) and manually curated. The Maximum-Likelihood phylogenetic tree was constructed using PhyML 3.0 with General Time Reversible (GTR) model and branch support computed by approximate likelihood ratio test (aLRT) method (Guindon et al. 2010).

### Genes annotation

The functional annotation of the genes was updated from the previous genome annotation (Madoui et al. 2017) using InterProScan v5.8-49.0 (Jones et al. 2014), BlastKOALA v2.1 (Kanehisa, Sato, and Morishima 2016) and by alignment on NCBI non-redundant protein database using Diamond (*Buchfink, Xie, and Huso*n 2015). Furthermore, a list of *O. nana* genes under natural selection in the Mediterranean Sea was added based on previous population genomic analysis (Arif et al. 2019). We further considered the annotation provided by either (i) Pfam (Finn et al. 2014) or SMART (Letunic and Bork 2018) protein domains, (ii) GO terms (molecular function, biological process or cellular component) (Ashburner et al. 2000) (iii) KEGG pathways (Kanehisa et al. 2012) and (iv) presence of locus under natural selection. These four gene features were used to identify specific enrichment in a given set of genes using a hypergeometric test that estimates the significance of the intersection between a specific gene list and one of the four global annotation lists.

### HMM search for LDPGs identification

From the InterProScan annotation of the *O. nana* proteome, 25 LNR domain sequences were detected (*p-value* ≤10-6), extracted and aligned with MAFFT using default parameters (Katoh and Standley 2013). A Hidden Markov Model (HMM) was generated from the aligned sequences using the ‘*hmmbuild*’ function of the HMMER tool version 3.1b1 (Eddy 2011). The *O. nana* proteome was scanned by ‘*hmmsearch*’ using the LNR HMM profile. Detected domains were considered as canonical LNR domains for *E-value, c-E-value* and *i-E-value* <10^−6^ and containing at least six cysteines or considered as LNR-like domains for *E-value, c-Evalue* and *i-Evalue* between 10^−6^ and 10^−1^ and containing at least four cysteines. A weblogo (Crooks et al. 2004) was generated to represent the conserved residues for the three LNRs of Notch protein, and the LNRs and LNR-like detected by HMM. The LNR and LNR-like domain-containing proteins constitute the LDPs final set used further. Deep-Loc (online execution) (Almagro Armenteros et al. 2017) was used to determinate the LDPs cellular localisation. To detect signal peptides and membrane protein topology, we used the online services of SignalP 5.0 (Almagro Armenteros et al. 2019) and TOPCONS (Tsirigos et al. 2015), respectively.

### Phylogeny tree of *O. nana* LNR domains

The *O. nana* nucleotide sequences of the LNR and LNR-like domains were aligned using MAFFT with default parameters. The Maximum-Likelihood phylogenetic tree was constructed using PhyML.3.0 with a model designed by the online execution of Smart Model Selection v.1.8.1 (Lefort, Longueville, and Gascuel 2017) and with branch supports computed by the aLRT method. The GTR model was used with an estimated discrete Gamma distribution (-a= 1.418) and a proportion of fixed invariable sites (−I = 0.25). The tree was visualised using MEGA-X (Kumar et al. 2018).

### Differential expression analysis

RNA-seq reads from the 20 libraries were mapped, independently, against the *O. nana* virtual cDNA, with ‘*bwa-mem*’ (v. 0.7.15-r1140) using default parameters (Li 2013) and read counts were extracted from the 20 BAM files with samtools (v. 1.4) (Li et al. 2009). Each reads set was validated by pairwise MA-plot to ensure a global representation of the *O. nana* transcriptome in each sample (Supplementary Notes S2). One nauplius sampled showing a biased read count distribution was discarded. Read counts from valid replicates were used as input data for the DESeq R package (Anders and Huber 2010) to identify differential gene expression between the five development stages through pairwise comparisons of each developmental stages. Genes having a Benjamini-Hochberg corrected p-value ≤ 0.05 in one of the pairwise comparisons were considered significantly differentially expressed. To identify stage-specific genes among these genes, we selected those being twice more expressed based on the normalised read count mean (log_2_(foldChange) > 1) in one development stage comparing to the four others. Up-regulated stage-specific genes were represented by a heatmap. The same method was used to determined down-regulated stage-specific genes (with log2(foldChange)<-1).

### Protein-protein interaction assays by yeast two-hybrid screening

Yeast two-hybrid experiments were performed using Matchmaker Gold Yeast Two-Hybrid System (Takara). The coding sequences of interest were first cloned into the entry vector pDONR/Zeo (ThermoFisher) and the correct ORF sequence verified by Sanger sequencing. To this aim, LDPGs were PCR-amplified with Gateway-compatible primers (Supplementary Notes S6) using cDNAs of pooled male individuals as template. In the case of secreted proteins, the amplified ORF lacked the signal peptide. Then, the cloned ORFs were reamplified by a two step-PCR protocol allowing the creation of a recombination cassette containing the ORF flanked by 40-nucleotide tails homologous to the ends of the pGBKT7 bait vector at the cloning site. Linearised bait vectors and ORF cassettes were co-transformed in Y2HGold yeast strain, and ORF cloning was obtained by homologous recombination directly in yeast. Y2H screening for potential interacting partners of the baits was performed against a cDNA library obtained from a pool of total mRNA of 100 *O. nana* male individuals constructed into the pGAD-AD prey vector.

Before screening, the self-activity of each bait clone was tested by mating with the Y187 strain harbouring an empty pGADT7-AD vector and then plating on SD/-His/-Leu/-Trp/ medium supplemented with 0, 1, 3, 5 or 10 mM 3-amino 1,2,4-triazole (3-AT). Each bait clone was then mated with the prey library containing approximately 4 × 10^6^ individual clones and plated on low-stringency agar plates (SD/-Trp/-Leu/-His/) supplemented with the optimal concentration of 3-AT based on the results of the self-activity test. To decrease the false positive rate, after five days of growth at 30°C, isolated colonies were spotted on high-stringency agar plates (SD/-Leu/-Trp/-Ade/-His) supplemented with 3-AT and allowed to grow another five days. Colony PCR on positive clones on this high stringency medium was performed with primers flanking the cDNA insert on the pGAD-AD vector, and PCR products were directly Sanger sequenced.

## Results

### *Oithona nana* female-biased sex-ratio

Between 2002 and 2017, 186 samples were collected in the Toulon Little Bay (figure 1.a), from which *O. nana* female and male adults were isolated. (figure 1.b). Across the fifteen years of observations, we noted a minimum male/female ratio in February (0.11), maxima in September, October and November (0.17) and a mean sex-ratio of 0.15 ± 0.11 all over the years (figure 1.c). This monitoring showed a relative stability along the year (ANOVA, P=0.87) but strongly biased sex-ratio toward females.

**Figure 1:**
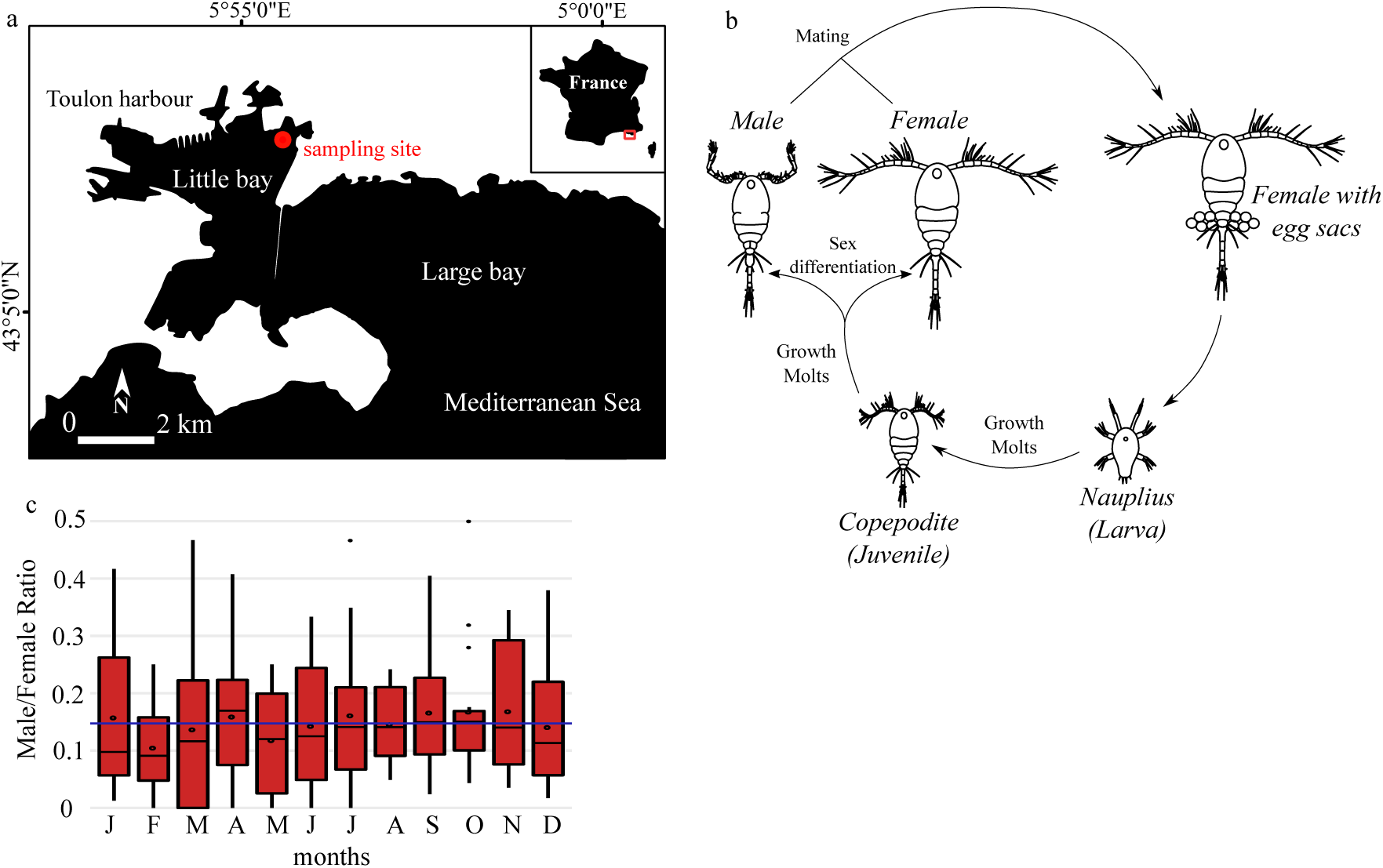
Life cycle and sex-ratio of the copepod *Oithona nana* in the Toulon Little Bay. **a.** Sampling station map in the Toulon Little Bay**. b**. The life cycle of *O. nana*. **c**. Sex ratio of *O. nana* time series in the Toulon Little Bay from 2002 to 2017. Black circles represent the mean by month. The blue line represents the fifteen-years mean (0.15).

### Male homogamety

To identify the most likely cause of this sex-ratio bias between environment sex determination (ESD) and higher male mortality, we used SD-pop on four individual transcriptomes of both sexes to determine the *O. nana* sexual system. According to SD-pop, the ZW model was preferred (lowest BIC) for *O. nana*. This result is unlikely to be due to chance, as for none of the runs on the 69 datasets for which the sex was permuted, the ZW model had the lowest BIC. Eleven genes had a posterior probability of being sex-linked in *O. nana* greater than 0.8. None of the SNPs in these genes showed the typical pattern of a fixed ZW SNP, i.e. the four females heterozygote and the four males homozygote (although, for some SNPs for which one individual was not genotyped, all four females were heterozygote, and all three genotyped males homozygote), indicating that the recombination suppression between the gametologs is recent, and that no or few mutations have gone to fixation independently in both gametolog copies. Annotation of these eleven genes shows that only one has homologs in other metazoans, ATP5H, that codes a subunit of mitochondrial ATP synthase (Supplementary Notes S8). Like in *Drosophila*, this gene is located in the nucleus (Liao et al. 2006).

### LNR domains burst in the *O. nana* proteome

To identify LDPs, we developed a HMM dedicated to *O. nana* LNR identification based on 31 conserved amino-acid residues. In the *O. nana* proteome, 178 LNR and LNR-like domains were detected and coded by 75 LDPGs, while a maximum of eight domains coded by six LDPGs was detected in the four other copepods (figure 2.b). Among the 178 *O. nana* domains, 22 were canonical LNR and 156 LNR-like domains (figure 2.c). By comparing the structure of Notch LNRs and LNR-like, we observed the loss of two cysteines (figure 2.c) in the LNR-like domains. Among the 75 LDPs, we identified nine different protein structure patterns (figure 2.d), including notably 47 LNR-only proteins, 12 trypsin-associated LDPs and eight metallopeptidase-associated LDPs. Overall, LDPs were predicted to contain a maximum of 5 LNR domains and 13 LNR-like domains.

**Figure 2:**
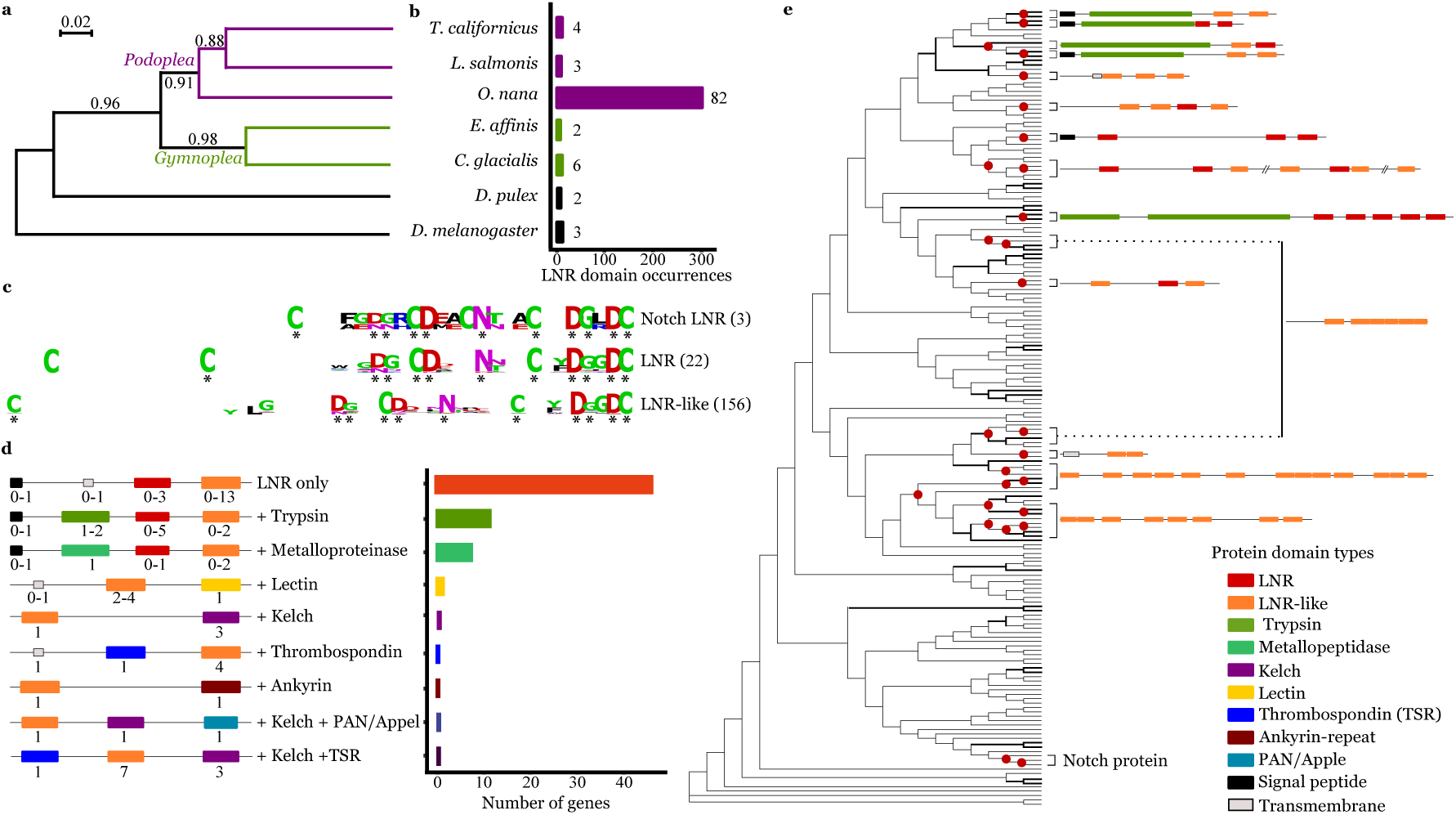
Lin-12 Notch Repeat (LNR) protein domain burst and high divergence with new domain associations in the *Oithona nana* proteome. **a**. Phylogeny of five copepod species and two other arthropod species based on 18S ribosomal sequences. The numbers at internal branches show the aLRT branch support. The scale bar represents the nucleotide substitution rate. **b**. LNR domain occurrences in seven Arthropoda proteomes detected by HMM. In front of each bar corresponds to the number of detected genes. **c**. Consensus sequences of the *O. nana* Notch LNR, LNR and LNR-like domains generated by WebLogo. The asterisks represent the conserved sites. **d**. Schemata of the *O. nana* LNR and LNR-like proteins structure. Numbers under each domain represent the possible occurrence range. The barplot represents the occurrence of the nine structures. **e**. Phylogenetic tree of the *O. nana* LNR and LNR-like domains. Bold branches have aLRT support ≥ 0.90. The red circles represent tandem duplication.

Forty-nine LDPs were predicted as secreted (eLDP), six membranous (mLDPs) and twenty intracellular (iLDPs) (Supplementary Notes S3). Among the iLDPs, two were associated with proteolytic domains, three associated with sugar-protein or protein-protein interaction domains (PAN/Appel, Lectin and Ankyrin) and 13 (65%) were LNR-only proteins. Among the eLDP, 18 (37%) contained proteolytic domains corresponding to a significant enrichment of proteolysis in eLDPs (*p-value*=2.13e-17); other eLDPs corresponded to LNR-only proteins (63%). The mLDPs were constituted of one Notch protein, two proteins with LNR domains associated with lectin or thrombospondin domains respectively, and three LNR-only proteins.

In the LNR and LNR-like domains phylogenetic tree bases on nucleic sequences (figure 2e), only 17% of the nodes had a support over 90%. Twenty-seven branch splits corresponded to tandem duplications involving 15 LDPGs, including Notch and a cluster of five trypsin-associated LDPGs codings three eLDPs and two iLDPs.

### *Oithona nana* male gene expression

Among the 15,399 genes predicted on the *O. nana* reference genome, 1,233 (~8%) were significantly differentially expressed in at least one of the five developmental stages. Among them, 619 genes were specifically up-regulated in one stage, with 53 genes up-regulated in eggs, 19 in nauplii, 75 in copepodids, 27 in adult females and 445 in adult males (figure 3.a). The male up-regulated genes were categorised based on their functional annotation (figure 3.b).

**Figure 3:**
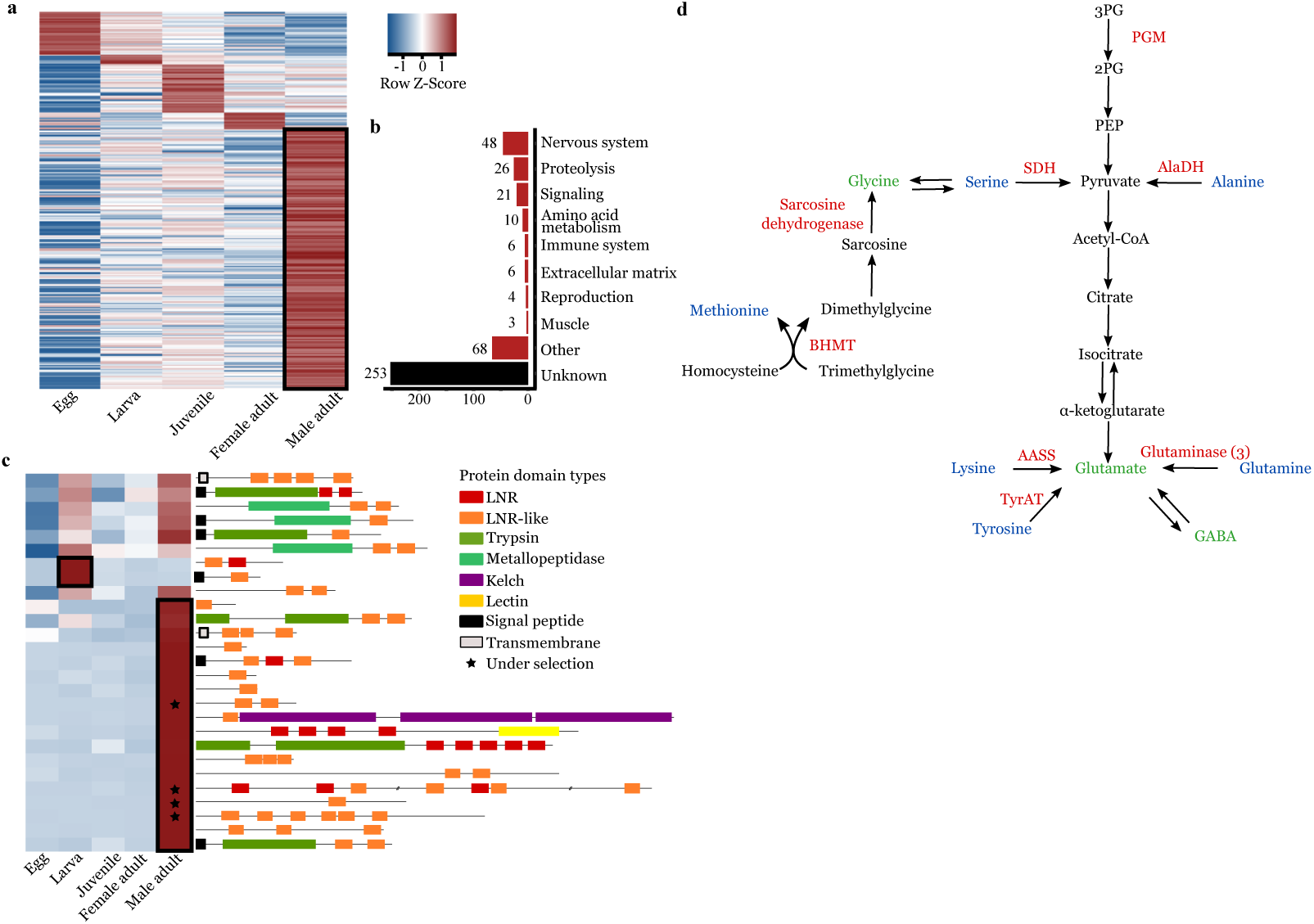
Differential expression analysis of the *Oithona nana* transcriptomes. **a**. Heatmap of the 1,233 significantly differentially expressed genes in at least one of the five developmental stages. **b**. Functional annotation distribution of 445 genes explicitly overexpressed in male adults. **c**. Heatmap of the 27 significantly differentially expressed LDPGs and their protein domains composition. **d**. Amino acid conversion to neurotransmitters in *O. nana* male. Over-expressed enzymes in males are in red, amino acids in blue and neurotransmitter amino acids in green.

#### Up-regulation of LNR-coding and proteolytic genes in adult male

The 1,233 differentially expressed genes contained 27 LDPGs (36% of total LDPGs) (figure 3.c). Over these 27 genes, 18 were specifically up-regulated in adult males producing a significant and robust enrichment of LDPGs in the adult males transcriptomes (fold > 8; *p-value*=2.95e-12) (figure 3.c). Among the 445 male-specific genes, 27 are predicted to play a role in proteolysis including 16 trypsins with three trypsin-associated LDPGs and showing significant enrichment of trypsin coding genes in males (*p-value*=1.73e-05), three metalloproteinases and five proteases inhibitors.

#### Up-regulation of nervous system associated genes in male adult

Forty-eight up-regulated genes in males are predicted functions in the nervous system (Supplementary Notes S4). These included 36 genes related to neuropeptides and hormones, through their metabolism (10 genes) with seven enzymes involved in neuropeptide maturation and one allatostatin, through their transport and release (9 genes), and through neuropeptide or hormone receptors (17 genes), seven of which are FMRFamide receptors. Six genes are predicted involved in the neuron polarisation, four in the axonal and dendrites organisation and growth guidance (including homologs to B4GAT1, futsch-like and zig-like genes), two in the development and maintenance of sensory and motor neurons (IMPL2, and DYF-5) and one in synapse formation (SYG-2).

#### Up-regulation of amino-acid conversion into neurotransmitters in male adults

We observed ten up-regulated genes in males predicted to play a role in amino acid metabolism (figure 3.b). This includes five enzymes that convert directly lysine, tyrosine and glutamine into glutamate through the activity of one α-aminoadipic semialdehyde synthase (AASS), one tyrosine aminotransferase (TyrAT) and three glutaminases, respectively. Three other enzymes might play a role in the formation of pyruvate: one alanine dehydrogenase (AlaDH), one serine dehydrogenase (SDH), and indirectly one phosphoglycerate mutase (PGM). Furthermore, two other enzymes might be involved in the formation of glycine, one sarcosine dehydrogenase (SARDH) and one betaine-homocysteine methyltransferase (BHMT) (figure 3.d).

#### Food uptake regulation in male adult

Three genes, which had predicted functions in food uptake regulation, showed specific patterns in male. These included the increase of the allatostatin-coding gene expression, a neuropeptide known in arthropods to reduce food uptake, but also three male under-expressed genes, a crustacean cardioactive peptide (CCAP), a neuropeptide that triggers digestive enzymes activation and the two bursicon protein subunits. These latter two hormones are known to be involved in intestinal and metabolic homeostasis.

### Protein-protein interaction involving LDPs and IGFBP

In order to further characterise the function of LDPGs, we studied potential proteins interactions by Yeast two-hybrid (Y2H) analysis (Supplementary Notes S5). To this aim, we selected eleven genes: seven male-overexpressed LDPGs, and four potential IGFBPs (Supplementary Notes S6).

We performed Y2H analysis by two different approaches (Supplementary Notes S5): the first was a matrix-based screen where potential binary interactions within candidate proteins were tested one-to-one. The second approach aimed to identify potential interactors in the entire *O. nana* proteome by a random library screen. This latter screening approach was more time-consuming and applied only to a subset of four genes (two LDPGs and two IGFBPs) used as baits against a Y2H library constructed from *O. nana* cDNAs.

Together, these two approaches allowed the reconstruction of a protein network containing 17 proteins including two LDPs and one IGFBP used as baits (figure 4.a), and 14 interacting partners of which six have an ortholog in other metazoans and five have no ortholog but at least one detected InterProScan domain (figure 4.b).

**Figure 4:**
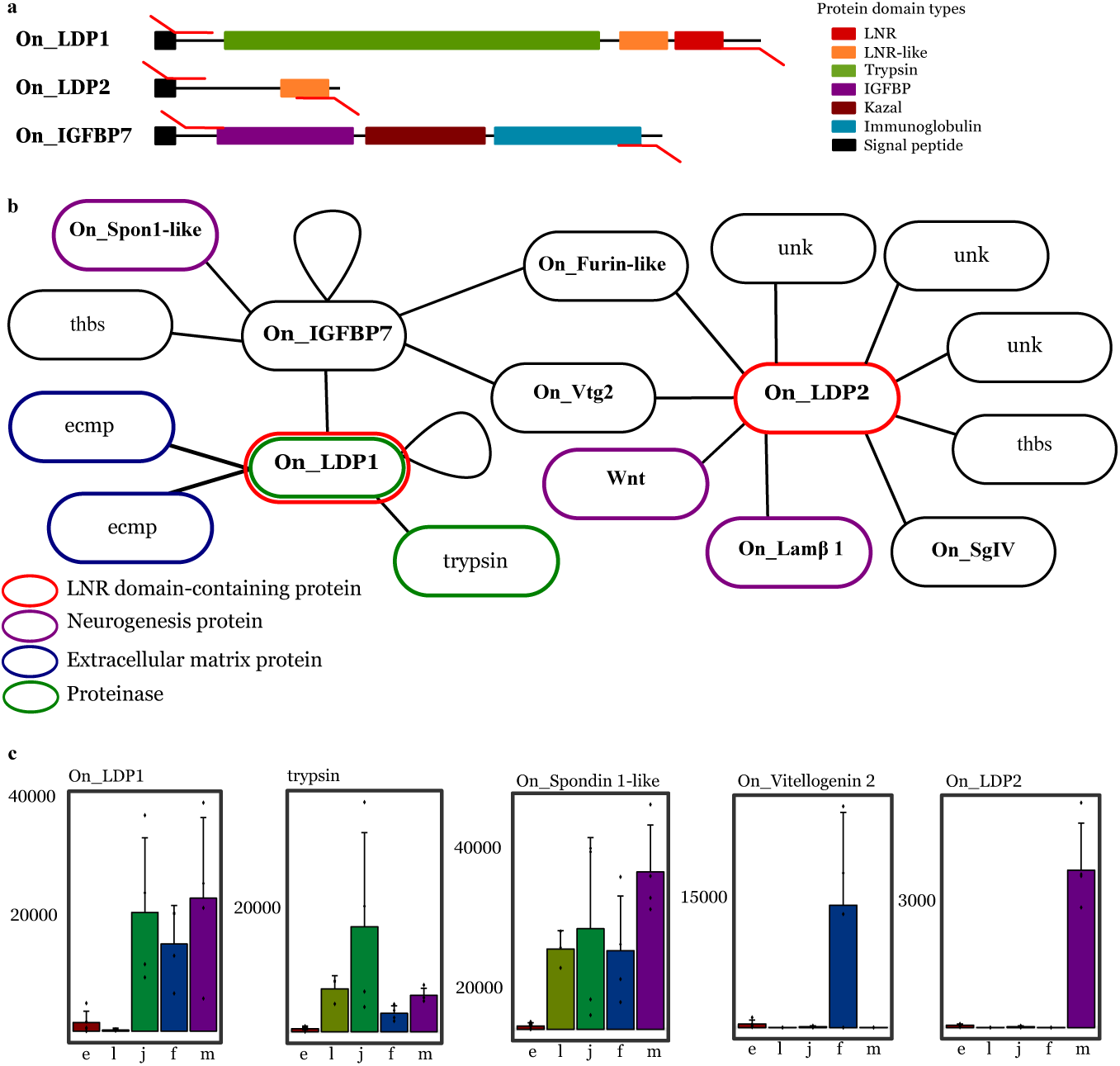
Protein-Protein Interaction of LNR-containing proteins in the *O. nana* male proteome. **a**. Structure and expression of the PPI candidates. The red arrows represent the PCR primers. **b**. PPI network of LDPGs obtained by Yeast two-hybrid assays. Lamβ 1: Laminin subunit beta 1; Spon1-like: Spondin1-like; Vtg2: Vitellogenin2; SgIV: Secretogranin IV; thbs: thrombospondins; unk: unknow. **c**. RPKM normalised expression for the five developmental stages. Only the five statistically differentially expressed genes are shown. From left to right: egg (e), larva (l), juvenile (j), female adult (f), male adult (m).

On_LDP1, an extracellular trypsin-containing LDP, formed a homodimer and interacted with a trypsin, two extracellular matrix (ECM) proteins and also an insulin-like growth factor binding protein (On_IGFBP7) that contains a trypsin inhibitor kazal domain. Based on its phylogeny, this protein is homolog to IGFBP7, also present in vertebrates (Supplementary Notes S7). On_IGFBP7 formed a homodimer and interacted with three other proteins: one spondin-1 like protein (On_Spon1-like) containing a kazal domain, one thrombospondin domain-containing protein and one vitellogenin 2-like protein (On_Vtg2). On_LDP2 is coded by a gene up-regulated in male (figure 4.c), detected under selection (Arif et al. 2019) and interacted with nine proteins: one vitellogenin 2-like protein (same interactant as On_IGFBP7), three uncharacterized proteins, one thrombospondin domain-containing protein (different than the On_IGFBP7 partner); one secretogranin V-like protein, one wnt-like protein, one laminin 1 subunit β and one a furin-like protein. No PPIs with IGF were detected, and no homolog of insulin-like androgenic gland hormone (Ventura, Rosen, and Sagi 2011) was found in the *O. nana* proteome.

## Discussion

### High mortality rate of *O. nana* males

Over 15 years of sampling, we observed a stable and strongly female-biased sex-ratio (~1:9) in the Toulon Little Bay. A similar observation was done in another *O. nana* population (Temperoni et al. 2011) and in other 132 *Oithonidae* populations (Kiørboe 2006). Two main causes could lead to these observations: a higher mortality of males or an environment-induced sex determination. As we showed that the *O. nana* sexual system is likely to be ZW, which would conduct to an expected 1:1 sex-ratio, the higher mortality rate of males seems more likely to explain our observations. These results are in accordance with the previously described risky behaviour of males, that is frequently in motion to find females and thus more vulnerable to predators than immobile females (Hirst et al. 2010).

### LDPs driven proteolysis in *O. nana* males

The explosion of LDPGs in the *O. nana* genome is unique in metazoan and is associated with the formation of new protein structures containing notably proteolytic domains. Owing to the LNR domain shortness (~40 amino acids) and the substantial polymorphism within the LNR domain sequences, the deep branches of the tree are weakly supported, and the evolutionary scenario of the domain burst remains hard to determine. However, the duplications of genes located in different scaffolds suggest post-duplication chromosomal rearrangements. Two previous studies on *O. nana* population genomics (Madoui et al. 2017; Arif et al. 2019) identified five LDPGs under natural selection with point mutations within an LNR domain. These results reinforce the idea of an ongoing evolution of these domains, forming new structures and thus allowing the emergence of new functions, especially in *O. nana* males according to the expression pattern of the LDPGs.

In metazoa, LNR domains are known to be involved in extracellular PPI (Boldt and Conover 2007) and cleavage site accessibility modulation (Sanchez-Irizarry et al. 2004). In *O. nana*, iLDPs are the most abundant type of LDPs. Their associations with other protein-binding domains like Kelsh, Ankyrin, PAN/Apple and thrombospondin repeat support a role of iLDPs in intracellular PPI. Half of the eLDPs are LNR-only and the other half is associated with peptidases (trypsin and metalloproteinase). From the PPI network, we showed that two eLDPs (On_LDP1 and On_LPD2) might interact with different types of extracellular proteins involved notably in tissue structure, energy storage and extracellular proteolysis. On the other hand, transcriptomic analyses showed an enrichment of trypsins in male adults (figure 3.c). The upregulation of allatostatin (Hergarden, Tayler, and Anderson 2012) and the downregulation of CCAP (Žitňan and Daubnerová 2016) and bursicon (Scopelliti et al. 2019) allow the male to reduce its food uptake. Taken together, this information supports a self-digestion of extracellular proteins in males driven by eLDPs and trypsin complexes that could act on proteolysis specificity modulation and/or on targeting/protecting specific extracellular proteins. This autolysis allows the release of amino acid and energy not supplied by feeding. Thus, the male adult autolysis could permit an expansion of mating period to increase its chances of mating (Heuschele and Kiørboe 2012) without motionless feeding phases. On the other hand, the deleterious effect of the autolysis on the organism supports a molecular-scale sacrificial behaviour in *O. nana* male.

### Neurotransmitter biosynthesis and nervous system development in *O. nana* male

From gene expression profile in *O. nana* male, we highlighted the direct and indirect conversion of four amino acids to glutamate, an excitatory neurotransmitter in arthropods, and the production of proteins composing the neurotransmitter vesicle transport which is consistent with the hyperactivity of the males during mate search. From the upregulation of neuronal developmental genes in male adult, and especially syg-2 and zig-8 normally expressed during the larval phase (Shen, Fetter, and Bargmann 2004), we infer an ongoing formation of new axons and/or dendrites and synapses in the male motor and/or sensory neurons. These results suggest a sexual dimorphism of the *O. nana* nervous system, as recently demonstrated in *Caenorhabditis elegans* (Cook et al. 2019).

Moreover, On_LDPG2, a male over-expressed gene and under natural selection in Mediterranean Sea populations, has been shown to interacts in yeast with two proteins involved in the nervous system development (On_Wnt and On_Lamβ1), notably in axon guidance (Zou 2004; Randlett et al. 2011). So, through its interactions, On_LDP2 may modulate neurogenesis in males and participate to the sexual dimorphism of the *O. nana* nervous system.

## Conclusion

In the Toulon Little Bay, *O. nana* presents a strong biased sex-ratio toward female while a ZW sexual determination system is favoured, which supports a higher mortality rate in male that can be explained by the sacrificial behaviour of males due to non-feeding, high motility and autolysis. The explosion of LDPGs in the *O. nana* genome seems to play an important role in the male-specific neurogenesis and autolysis. However, more investigation should be undergone to identify which part of the nervous system is developing and which tissues are lysed. To our knowledge, sacrificial behaviour was only observed in semelparous animals. Thus, the sacrificial behaviour supported at the molecular level in *O. nana* by autolysis may represent the first example of sacrificial behaviour in semelparous animals. Subtle trade-offs and molecular control mechanisms for the autolysis of specific tissues may occur to adjust the autolysis to the lifespan of the male allowing it to reproduce several times. The low mortality rate of the motionless females and the male sacrificial behaviour could be one of the main factors of the ecological success of *O. nana (Razouls et al. 2019)*.

## Acknowledgements

We acknowledge the support of the Genoscope-CEA, France Génomique (ANR-10-INBS-09) and the French Ministry of Research.

## Data availability

The *O. nana* RNA-seq data are available at ENA (*Supplementary Notes* S1)

## Authors’ contribution

KS, JP and JLJ collected the samples; JLJ generated the sex-ratio data; KS, KL, MAM, EP and JP generated the molecular data; JK performed sexual system analysis; KS, AA, LB, SK, NM, and CO performed the yeast two-hybrid analysis; AA designed the yeast two-hybrid method; KS and MAM performed the analyses; KS, AA and MAM wrote to the manuscript; MAM supervised the study.

## Figure Legends

Supplementary Notes S1: **Transcriptomic data.** RNA-seq quality metrics of the 20 samples. One nauplii sample was discarded after MA plots pairwise comparisons analysis (see Supplementary Notes 2).

Supplementary Notes S2: **Transcriptomes quality**. MA plots pairwise comparisons between the five developmental stages. One nauplius sampled showing a biased read count distribution was discarded.

Supplementary Notes S3: **Structure and localisation of the *Oithona nana* LDPs**. *e, i* and *m* correspond to extracellular, intracellular and membranous respectively

Supplementary Notes S4: **Functional annotation of *O. nana* genes over-expressed in male**.

Supplementary Notes S5: **Experimental design of the protein interaction (PPI) analysis**.

Supplementary Notes S6: **Primers used for PCR amplification**. The first table contains the primers used for bait amplification; the second contains the prey ones.

Supplementary Notes S7: **Phylogenetic tree of On_IGFBP7**. The numbers at internal branches show the bootstrap branch support (100). We used NCBI data from *Mus muculus* (mouse), *Danio rerio* (danre) and *Bos taurus* (bovin).

Supplementary Notes S8. **Gene annotation of the sex-determination system associated genes of *Oithona nana***

