## Supplementary Notes for "Proteolysis and neurogenesis modulated by LNR domain proteins explosion support male differentiation in the crustacean *Oithona nana*"

Supplementary data

### Supplementary Notes S1**:** **Transcriptomic data.** RNA-seq quality metrics of the 20 samples. One nauplii sample was discarded after MA plots pairwise comparisons analysis (see Sup note 2).

| Accession Numbers  (PRJEB34229) |  | Biological replicate | # Seq clean | # Base clean | Average size of merged reads (base) |
| --- | --- | --- | --- | --- | --- |
| ERX3525067 | Male | 1 | 17,539,769 | 4,726,772,098 | 155 |
| ERX3525068 |  | 2 | 13,355,099 | 3,594,123,134 | 155 |
| ERX3525069 |  | 3 | 13,327,828 | 3,697,975,710 | 164 |
| ERX3525070 |  | 4 | 10,181,747 | 2,833,631,554 | 164 |
| ERX3525072 | Female | 1 | 13,443,737 | 3,784,911,561 | 169 |
| ERX3525073 |  | 2 | 17,565,613 | 4,945,908,456 | 170 |
| ERX3525074 |  | 3 | 15,129,175 | 4,254,430,406 | 170 |
| ERX3525071 |  | 4 | 13,152,019 | 3,685,066,788 | 167 |
| ERX3525076 | Copepodid | 1 | 9,737,697 | 2,683,086,720 | 159 |
| ERX3525077 |  | 2 | 10,954,752 | 3,090,141,346 | 170 |
| ERX3525078 |  | 3 | 13,385,311 | 3,732,623,083 | 166 |
| ERX3525075 |  | 4 | 17,758,141 | 4,731,918,511 | 152 |
| ERX3525079 | Nauplius | 1 | 14,280,023 | 3,868,981,343 | 158 |
| ERX3525080 |  | 2 | 13,034,144 | 3,546,435,298 | 159 |
| ERX3525081 |  | 3 | 11,718,354 | 3,291,309,412 | 169 |
| ERX3525082 | Egg | 1 | 16,670,807 | 4,457,103,808 | 153 |
| ERX3525083 |  | 2 | 14,351,720 | 3,906,557,216 | 159 |
| ERX3525084 |  | 3 | 12,333,551 | 3,284,934,971 | 152 |
| ERX3525085 |  | 4 | 17,427,615 | 4,468,263,185 | 142 |
| ERX3525086 |  | 5 | 16,156,218 | 4,271,517,322 | 150 |

### Supplementary Notes S2: **Transcriptomes quality.** MA plots pairwise comparisons between the five developmental stages. One nauplius sampled showing a biased read count distribution was discarded.

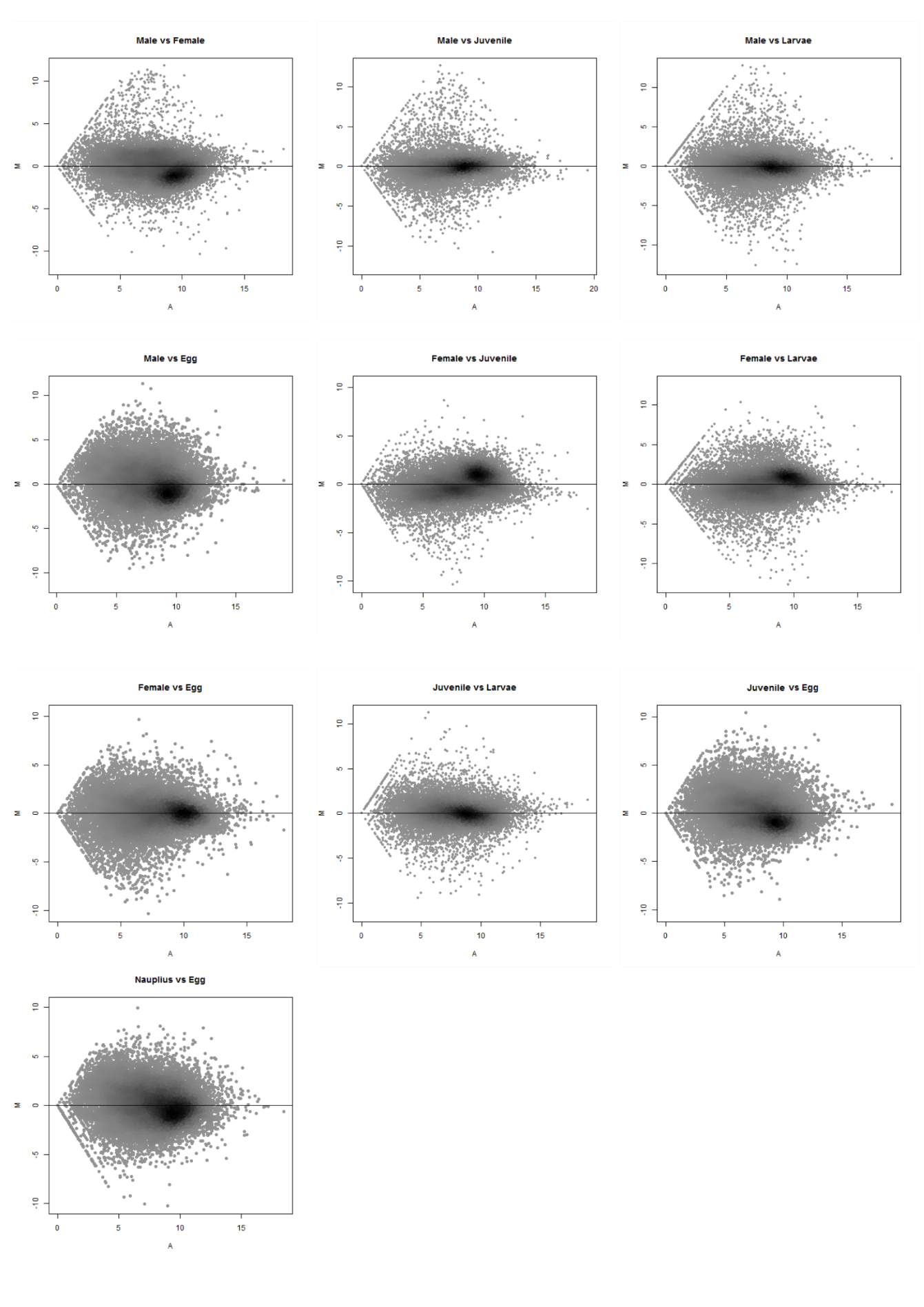

### Supplementary Notes S3: **Structure and localisation of the *Oithona nana* LDPs.** *e, i* and *m* correspond to extracellular, intracellular and membranous respectively.

| Protein name | Protein size (aa) | #Total LNR domain (#canonical LNR - # LNR-like) | Other InterPro domains | Localisation |
| --- | --- | --- | --- | --- |
| 100:211670..212713 | 328 | 3 (1-2) | - | e |
| 11.480.1:721370..723090 | 531 | 1 (0-1) | Metallo-peptidase family M12 | e |
| 12.169.2:773247..776065 | 808 | 5 (5-0) | Trypsin | e |
| 12.175.1 | 386 | 1 (0-1) | Trypsin | e |
| 12.182.1 | 445 | 2 (0-2) | Trypsin | e |
| 12.185.1 | 357 | 1 (0-1) | Trypsin | e |
| 123:42215..42819 | 181 | 2 (0-2) | - | e |
| 1239:190..1043 | 228 | 2 (0-2) | - | e |
| 14.373.1:814017..816142 | 226 | 2 (0-2) | - | e |
| 1531:53..2073 | 222 | 3 (0-3) | - | e |
| 1646:48..560 | 147 | 1 (0-1) | - | e |
| 174:49279..49938 | 219 | 3 (0-3) | - | e |
| 1781:78..483 | 113 | 1 (1-0) | Metallo-peptidase family M12B | e |
| 18.220.1:802020..803354 | 420 | 1 (0-1) | Trypsin | e |
| 188.202.1:42571..49247 | 547 | 2 (2-0) | - | e |
| 19.334.1 | 365 | 4 (1-3) | - | e |
| 2.84.1 | 195 | 2 (1-1) | Trypsin | e |
| 20.537.1 | 353 | 2 (1-1) | - | e |
| 2085.199.1 | 687 | 6 (0-6) | - | e |
| 21.77.1:327349..328866 | 458 | 2 (1-1) | Trypsin | e |
| 2201:1341..2883 | 426 | 3 (0-3) | - | e |
| On_LDP1 | 378 | 2 (1-1) | Trypsin | e |
| 27:531099..531903 | 248 | 2 (0-2) | - | e |
| 32.270.1:20364..22049 | 561 | 2 (0-2) | - | e |
| 3256:80..502 | 140 | 1 (0-1) | - | e |
| 37:34427..34747 | 107 | 1 (0-1) | - | e |
| 3704.33.1 | 266 | 1 (0-1) | Metallo-peptidase family M12 | e |
| 3765:1004..2202 | 383 | 1 (0-1) | - | e |
| 38:505765..507559 | 438 | 5 (0-5) | - | e |
| 3888:228..1050 | 202 | 2 (1-1) | - | e |
| 4.222.1 | 525 | 2 (0-2) | Metallo-peptidase family M12 | e |
| 4.25.1 | 90 | 1 (0-1) | - | e |
| 41:85441..86498 | 276 | 2 (0-2) | - | e |
| 42.291.1 | 122 | 1 (0-1) | - | e |
| 45.412.1 | 262 | 1 (0-1) | Trypsin | e |
| 45.424.1 | 139 | 1 (0-1) | - | e |
| 4546:128..817 | 229 | 3 (0-3) | - | e |
| 48.267.1:6982..8492 | 461 | 2 (0-2) | Trypsin | e |
| 48.344.1 | 525 | 2 (1-1) | Metallo-peptidase family M12 | e |
| 54.120.2:255915..258850 | 940 | 7 (0-7) | - | e |
| 55:11070..12199 | 357 | 4 (0-4) | - | e |
| 57.114.1 | 159 | 2 (1-1) | - | e |
| 6.168.1:380139..381062 | 227 | 1 (0-1) | Metallo-peptidase family M12B | e |
| 68.40.1 | 493 | 1 (0-1) | Metallo-peptidase family M12 | e |
| 821:3497..4457 | 291 | 6 (0-6) | - | e |
| On_LDP2 | 116 | 1 (0-1) | - | e |
| 92.33.1:248003..248641 | 213 | 2 (0-2) | - | e |
| 98.272.1 | 74 | 1 (1-0) | - | e |
| 99.319.1:158136..159228 | 344 | 1 (0-1) | Trypsin | e |
| 12.180.1 | 489 | 2 (0-2) | Trypsin | i |
| 126.234.1 | 823 | 2 (0-2) | - | i |
| 14:255249..255707 | 252 | 1 (0-1) | - | i |
| 15.286.1 | 207 | 2 (0-2) | - | i |
| 1807:1535..3342 | 517 | 1 (0-1) | - | i |
| 188:49497..50375 | 177 | 1 (1-0) | - | i |
| 1921.111.1 | 137 | 1 (0-1) | - | i |
| 2.15.1:36291..38976 | 767 | 13 (0-13) | - | i |
| 2.380.1:1294082..1295071 | 329 | 2 (0-2) | - | i |
| 20.536.3 | 1083 | 1 (0-1) | Kelch motif | i |
| 2184:300..2200 | 310 | 1 (0-1) | Ankyrin repeat-containing domain superfamily | i |
| 2277:24..1778 | 518 | 8 (0-8) | - | i |
| 23.148.2 | 460 | 2 (0-2) | Metallo-peptidase family M12 | i |
| 243.135.1 | 316 | 2 (0-2) | - | i |
| 284.43.1 | 132 | 1 (0-1) | - | i |
| 468:255..2448 | 602 | 1 (0-1) | PAN/Apple domain ; Kelch motif | i |
| 541.168.1 | 1564 | 7 (3-4) | - | i |
| 556:85..5023 | 1494 | 6 (0-6) | TSP1 ; Kelch-typ_b-propeller | i |
| 60:181372..182580 | 402 | 1 (0-1) | - | i |
| 94.274.1:231104..233771 | 867 | 4 (0-4) | Lectin | i |
| 147.226.2 | 266 | 3 (0-3) | - | m |
| 36:371592..372773 | 336 | 1 (0-1) | - | m |
| On_Notch | 2147 | 3(3-0) | Notch protein | m |
| 59:239420..240555 | 358 | 2 (0-2) | - | m |
| 76.32.1:162363..164050 | 435 | 4 (0-4) | Thrombospondin type-1 (TSP1) repeat superfamily | m |
| 96.130.1:322..1331 | 262 | 2 (0-2) | Lectin | m |

### Supplementary Notes S4: **Functional annotation of O. *nana* genes over-expressed in male.**

| **Protein functional groups** | **Protein** | **Number of male-specific genes** |
| --- | --- | --- |
|  | Allatostatin precursor protein | 1 |
|  | Carboxypeptidase E | 1 |
| **Neuropeptide and** | Furin-like protease | 1 |
| **hormone metabolism** | Neuroendocrine convertase 2 | 1 |
|  | Peptidyl-alpha-hydroxyglycine alpha-amidating lyase | 1 |
|  | Peptidylglycine alpha-hydroxylating monooxygenase-like | 1 |
|  | ITG-like peptide | 1 |
|  | Sulfoacetaldehyde acetyltransferase | 1 |
|  | ***Total*** | ***8*** |
|  | Bestrophin 1 | 1 |
|  | Excitatory amino acid transporter 1 | 1 |
| **Neuropeptide and** | Multiple C2 and transmembrane domain-containing protein | 1 |
| **hormone transport** | Sodium- and chloride-dependent GABA transporter 1-like | 1 |
| **and release** | Synaptic vesicular amine transporter | 1 |
|  | Synaptobrevin | 1 |
|  | Synaptotagmin | 1 |
|  | Synaptosomal-associated protein 25-like | 1 |
|  | Neuronal calcium sensor-like | 1 |
|  | ***Total*** | ***9*** |
|  | FMRFamide receptor | 6 |
|  | Muscarinic acetylcholine receptor | 1 |
|  | GABA receptor subunit | 2 |
|  | Glycine receptor subunit | 2 |
| **Neuropeptide and** | Ionotropic glutamate receptor subunit | 2 |
| **hormone receptors** | Capa receptor | 1 |
|  | Octopamine receptor subunit | 1 |
|  | Corticotropin hormone receptor | 1 |
|  | Acetylcholine receptor subunit | 1 |
|  | ***Total*** | ***17*** |
|  | Kv channel-interacting protein 1-like | 2 |
| **Neuron polarization** | Potassium voltage-gated channel, Shab-related subfamily | 2 |
| **Current propagation** | Transient receptor potential protein | 1 |
| **and ion channel** | Two pore potassium channel protein sup-9 | 1 |
|  | ***Total*** | ***6*** |
|  | Beta-1,4-glucuronyltransferase 1 | 1 |
| **Neuron development** | Microtubule-associated protein futsch-like | 1 |
| **Synapse assembly** | Serine/threonine-protein kinase dyf-5 ou ICK | 1 |
|  | Neural/ectodermal development factor IMP-L2-like | 1 |
|  | zwei Ig domain protein | 2 |
|  | copine-like | 1 |
|  | synaptogenesis syg-2 | 1 |
|  | ***Total*** | ***8*** |
| ***Total*** |  | ***48*** |

### Supplementary Notes S5: **Experimental design of the protein interaction (PPI) analysis.**

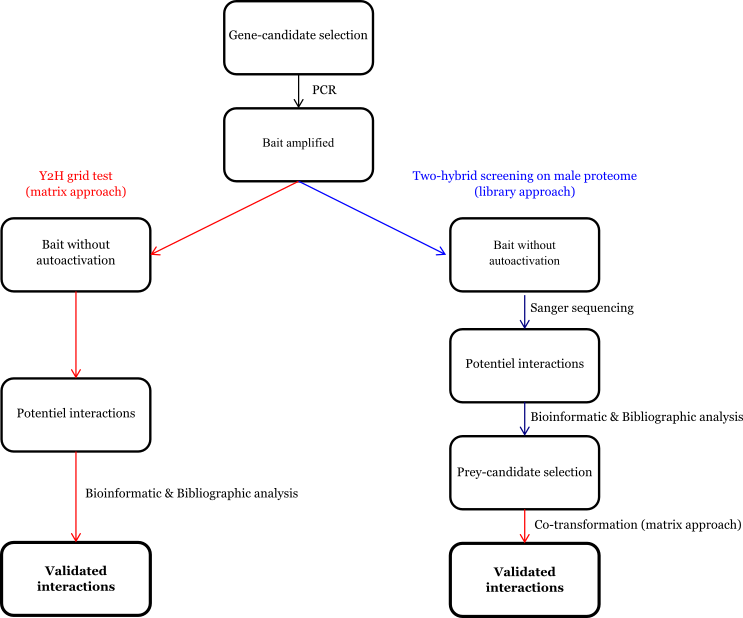

Supplementary Notes S6: P**rimers used for PCR amplification.** The first table contains the primers used for bait amplification; the second contains the prey ones.

| Gene | Elongation direction | Primer |
| --- | --- | --- |
| On_LDPG1 | Forward | ATGGCCATGGAGGCCGAATTCCTTCCAGGATTTGTTGGAAACTTG |
|  | Reverse | CCGCTGCAGGTCGACGGATCCGCGAATACATTGGCAAATGGTACA |
| On_LDPG2 | Forward | ATGGCCATGGAGGCCGAATTCCCCTATTGCCAAGCCTGTGGT |
|  | Reverse | CCGCTGCAGGTCGACGGATCCGGTTTGAGGATACCACCGATAATC |
| On_IGFBP7 | Forward | ATGGCCATGGAGGCCGAATTCCAACGTCGTCCACATGCCCCAAAT |
|  | Reverse | CCGCTGCAGGTCGACGGATCCCAATTCTTTACTATAAACGCCCAC |
| 4.25.1 | Forward | ATGGCCATGGAGGCCGAATTCACTAAAATGATTCTATCAGATCGC |
|  | Reverse | CCGCTGCAGGTCGACGGATCCTTCTTCTAGAATGATTTTATCATT |
| 19.110.1 | Forward | ATGGCCATGGAGGCCGAATTCATGTCAGTATCATATTTTAAAATC |
|  | Reverse | CCGCTGCAGGTCGACGGATCCGCAGTCTAAACAATGATCAATTAC |
| 49.77.3 | Forward | ATGGCCATGGAGGCCGAATTCCCACAATACTTTCAACTTGGAAAC |
|  | Reverse | CCGCTGCAGGTCGACGGATCCGAACTGGAAAATGTGGGGAATGAG |
| 45.413.1 | Forward | ATGGCCATGGAGGCCGAATTCTGTGTAGGCTGCGGAGAATACCCA |
|  | Reverse | CCGCTGCAGGTCGACGGATCCTTGACCGGATGTCGGAGGTGGAGG |
| 2085.199.1 | Forward | ATGGCCATGGAGGCCGAATTCTTAACCACCACCCCTATTGATCTA |
|  | Reverse | CCGCTGCAGGTCGACGGATCCCCGTGGCAATGTGCTAAATTGAGA |
| 12.180.1 | Forward | ATGGCCATGGAGGCCGAATTCTTGCCTTTCACTGTTTCCGACAAT |
|  | Reverse | CCGCTGCAGGTCGACGGATCCCCGTGGCAATGTGCTAAATTGAGA |
| 94.274.1 | Forward | ATGGCCATGGAGGCCGAATTCGATGATGGAGGATGTGCTTTTAGG |
|  | Reverse | CCGCTGCAGGTCGACGGATCCAGATGATCCTGTAGTGGTTGCGGT |
| 1921.111.1 | Forward | ATGGCCATGGAGGCCGAATTCGTACCCGAGTTTCAATTTGATGGA |
|  | Reverse | CCGCTGCAGGTCGACGGATCCATCAGGATGAAAGGTGCAGTCTTG |

| Gene | Elongation direction | Primer |
| --- | --- | --- |
| On_LDPG1 | Forward | GCCATGGAGGCCAGTGAATTCCTTCCAGGATTTGTTGGAAACTTG |
|  | Reverse | CAGCTCGAGCTCGATGGATCCGCGAATACATTGGCAAATGGTACA |
| On_LDPG2 | Forward | GCCATGGAGGCCAGTGAATTCTTTGAACTTTGCTTTTATGATGGA |
|  | Reverse | CAGCTCGAGCTCGATGGATCCGCAGTCTAAACAATGATCAATTAC |
| On_IGFBP7 | Forward | GCCATGGAGGCCAGTGAATTCCAACGTCGTCCACATGCCCCAAAT |
|  | Reverse | CAGCTCGAGCTCGATGGATCCCAATTCTTTACTATAAACGCCCAC |
| 4.25.1 | Forward | GCCATGGAGGCCAGTGAATTCACTAAAATGATTCTATCAGATCGC |
|  | Reverse | CAGCTCGAGCTCGATGGATCCTTCTTCTAGAATGATTTTATCATT |
| 19.110.1 | Forward | GCCATGGAGGCCAGTGAATTCACTAAAATGATTCTATCAGATCGC |
|  | Reverse | CAGCTCGAGCTCGATGGATCCGAAATCAAAAGGGTTTGGCACATT |
| 49.77.3 | Forward | GCCATGGAGGCCAGTGAATTCCCACAATACTTTCAACTTGGAAAC |
|  | Reverse | CAGCTCGAGCTCGATGGATCCGAACTGGAAAATGTGGGGAATGAG |
| 45.413.1 | Forward | GCCATGGAGGCCAGTGAATTCATGGACTTCAATTCCATTTATGAA |
|  | Reverse | CAGCTCGAGCTCGATGGATCCTTGACCGGATGTCGGAGGTGGAGG |
| 2085.199.1 | Forward | GCCATGGAGGCCAGTGAATTCTTAACCACCACCCCTATTGATCTA |
|  | Reverse | CAGCTCGAGCTCGATGGATCCCCGTGGCAATGTGCTAAATTGAGA |
| 12.180.1 | Forward | GCCATGGAGGCCAGTGAATTCTTGCCTTTCACTGTTTCCGACAAT |
|  | Reverse | CAGCTCGAGCTCGATGGATCCAGTTACAATGCATTCACAAACTTT |
| 94.274.1 | Forward | GCCATGGAGGCCAGTGAATTCGATGATGGAGGATGTGCTTTTAGG |
|  | Reverse | CAGCTCGAGCTCGATGGATCCAGATGATCCTGTAGTGGTTGCGGT |
| 1921.111.1 | Forward | GCCATGGAGGCCAGTGAATTCGTACCCGAGTTTCAATTTGATGGA |
|  | Reverse | CAGCTCGAGCTCGATGGATCCATCAGGATGAAAGGTGCAGTCTTG |

### Supplementary Notes S7**: Phylogenetic tree of On_IGFBP7.** The numbers at internal branches show the bootstrap branch support (100). We used NCBI data from *Mus muculus* (mouse), *Danio rerio* (danre) and *Bos taurus* (bovin).

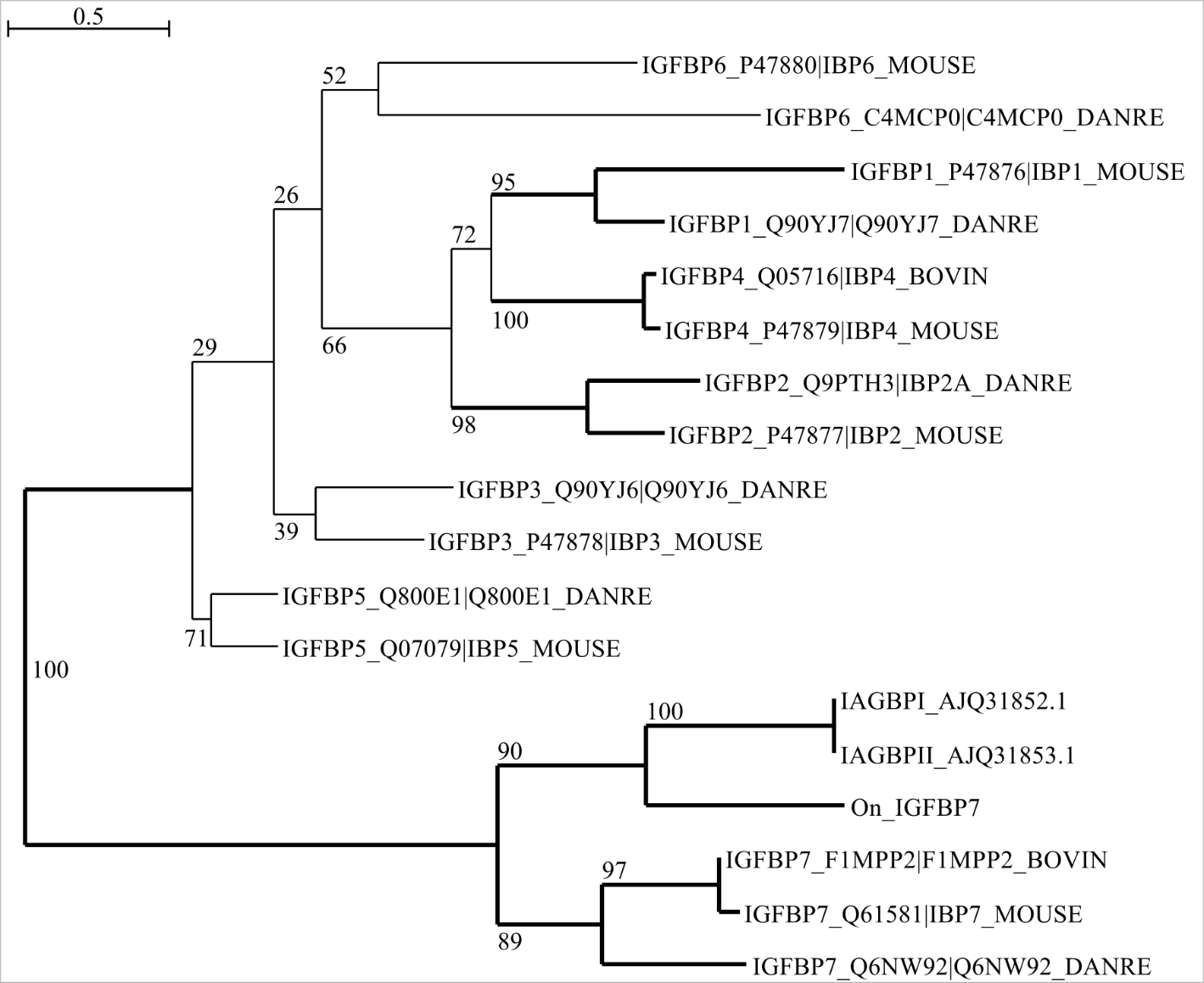

### Supplementary Notes S**8**. **Gene annotation of the sex-determination system associated genes of *Oithona nana***

| Gene | Diamond blast NR | InterProScan domains | RPKM average Female | RPKM average Male |
| --- | --- | --- | --- | --- |
| 7.623.1 | No hits |  | ~5 | ~24 |
| 34.133.1 | No hits |  | ~25 | ~80 |
| 39.231.1 | No hits |  | ~50 | ~20 |
| 87.175.1 | No hits |  | ~60 | ~110 |
| 45.363.1 | No hits | PAN domain | ~100 | ~10 |
| 25.213.1 | No hits |  | ~475 | ~10 |
| 664.74.1 | No hits |  | ~500 | ~120 |
| 27.64.1 | No hits |  | ~600 | ~250 |
| 41.148.1 | No hits |  | ~1200 | ~600 |
| 119.249.1 | putative protein LLP homolog isoform X1 [Acartia pacifica] | Learning-associated protein | ~1800 | ~800 |
| On_ATP5H | ATP synthase subunit d, mitochondrial-like [Eurytemora affinis] | ATP synthase D chain, mitochondrial (ATP5H) | ~7500 | ~21000 |
